# WOX11-mediated cell size control in Arabidopsis attenuates fecundity of endoparasitic cyst nematodes

**DOI:** 10.1101/2023.10.27.564344

**Authors:** Nina Guarneri, Jaap-Jan Willig, Viola Willemsen, Aska Goverse, Mark G. Sterken, Pieter Nibbering, Jose L. Lozano-Torres, Geert Smant

**Affiliations:** Laboratory of Nematology, Wageningen University & Research, 6708 PB Wageningen, the Netherlands; Laboratory of Cell and Developmental Biology, Cluster of Plant Developmental Biology, Wageningen University & Research, 6708 PB Wageningen, the Netherlands

**Author notes:** Author for correspondence: Geert Smant.

**Keywords:** cyst nematodes, WOX11, cell size, syncytium hypertrophy, female fecundity, cell wall plasticity, ROS

## Abstract

- Cyst nematodes establish permanent feeding structures called syncytia inside host root vasculature, disrupting the flow of water and minerals. In response, plants form WOX11-mediated adventitious lateral roots at nematode infection sites. WOX11-adventitious lateral rooting modulates tolerance to nematode infections, however, whether this also benefits nematode parasitism remains unknown.
- Here, we report on bioassays using a *35S::WOX11-SRDX* transcriptional repressor mutant to investigate whether WOX11-adventitious lateral rooting promotes syncytium development and thereby female fecundity. Moreover, we chemically inhibited cellulose biosynthesis to verify if WOX11 directly modulates cell wall plasticity in syncytia. Finally, we performed histochemical analyses to test if WOX11 mediates syncytial cell wall plasticity via reactive oxygen species (ROS).
- Repression of WOX11-mediated transcription specifically enhanced the radial expansion of syncytial elements, increasing both syncytium size and female offspring. The enhanced syncytial hypertrophy observed in the *35S::WOX11-SRDX* mutant could be phenocopied by chemical inhibition of cellulose biosynthesis and was associated with elevated levels of ROS at nematode infection sites.
- We therefore conclude that WOX11 restricts radial expansion of nematode feeding structures and female fecundity, likely by modulating ROS-mediated cell wall plasticity mechanisms. Remarkably, this novel role of WOX11 in plant cell size control is independent of WOX11-adventitious rooting underlying disease tolerance.

## Introduction

Biotic stress by endoparasitic cyst nematodes disrupts plant growth by altering the flow of water and minerals through destructive migration within the roots and feeding (Trudgill *et al*., 1975, Grundler and Hofmann, 2011, Rodiuc *et al*., 2014, Levin *et al*., 2021). In response to nematode infection, plants trigger a damage signaling pathway mediated by *WUSCHEL-RELATED HOMEOBOX11 (WOX11)* that leads to the formation of adventitious lateral roots at nematode infection sites (Guarneri *et al*., 2023, Willig *et al*., 2023). WOX11-adventitious lateral roots compensate for the inhibition of primary root growth caused by cyst nematode infection and contribute to better maintenance of aboveground plant development and growth (Willig *et al*., 2023). Thus, WOX11 modulates plant tolerance to cyst nematode infections (Willig *et al*., 2023). Furthermore, WOX11 reduces Arabidopsis susceptibility to cyst nematode penetration (Willig *et al*., 2023). However, whether WOX11 may affect cyst nematode feeding and thereby female fecundity remains unknown.

Cyst nematode females require a permanent feeding structure to reach the adult stage and produce eggs. Hereto, upon host penetration, the infective second-stage juveniles (J2s) insert their needle-like oral stylet into a cell of the plant vascular cylinder and secrete effector proteins. As a result, this cell undergoes a cascade of structural changes and fuses with hundreds of neighboring cells by partial cell wall dissolution to form the so-called syncytium. During this process, activation of the endocycle increases the DNA content, while syncytial elements expand radially by hypertrophy (Golinowski *et al*., 1996). Nematode juveniles feed on syncytia in cycles of continuous ingestion and resting periods, and molt three times until they become adult females and males (Muller *et al*., 1981). While males stop feeding after the third larval stage and are associated with relatively small syncytia, females ingest food until the adult stage and have large, hypertrophied syncytia (Muller *et al*., 1981, Hofmann and Grundler, 2006). Syncytial hypertrophy positively correlates with female size, which can be used as an indicator of female fecundity (Urwin *et al*., 1997, Goverse *et al*., 2000, Li *et al*., 2004, Siddique *et al*., 2012, Ali *et al*., 2013, Siddique *et al*., 2014, Siddique *et al*., 2015, Chopra *et al*., 2021, Siddique *et al*., 2022).

The hypertrophy of female-associated syncytia is thought to depend on the uptake of assimilates, such as sucrose from the phloem (Hofmann and Grundler, 2006). Initially, syncytia are symplastically isolated from surrounding host tissues and sucrose is taken up from phloem companion cells via active transport. Later, when syncytia have reached their maximum expansion, the opening of secondary plasmodesmata allows the passive transport of sucrose from the phloem sieve elements (Hofmann *et al*., 2007). Increased osmolarity due to the high sucrose concentration causes the passive inflow of water from the xylem, elevating turgor pressure in the syncytia (Böckenhoff and Grundler, 1994). High turgor pressure poses tensile stress on plant cell walls and is thought to drive syncytial hypertrophy (Hofmann and Grundler, 2006, Cosgrove, 2022). At the same time, modifications in the syncytial cell wall composition likely provide mechanical strength to withstand the turgor pressure, while allowing syncytial elements to expand and thus accommodate the periodic demands imposed by nematode feeding (Zhang *et al*., 2017). Indeed, silencing a cell-wall modifying enzyme involved in cellular hypertrophy compromised female fecundity in potato (Catalá *et al*., 2000, Karczmarek *et al*., 2008).

The presence of WOX11-mediated adventitious lateral roots at nematode infection sites might interfere with the flow of assimilates and water towards the syncytium, with possible consequences on syncytium hypertrophy and female fecundity. Indeed, similarly to syncytia, adventitious lateral roots constitute a sink of assimilates for the plant (Hofmann *et al*., 2007, Stitz *et al*., 2023). As such, adventitious lateral roots may compete with syncytia for the uptake of sucrose. Moreover, root branching depends on the activation of glycolysis in the roots, which increases the demand for shoot-derived carbon sources (Stitz *et al*., 2023). Consequently, adventitious lateral root formation may benefit syncytia by enhancing the overall availability of sucrose in the roots. Besides, mature adventitious lateral roots likely increase water flux towards the syncytium (Levin *et al*., 2020), thereby promoting turgor-driven syncytial hypertrophy. As a result, larger, hypertrophied syncytia may accumulate higher amounts of sucrose, thus better supporting nematode feeding. Therefore, we hypothesize that WOX11 affects syncytium hypertrophy and thus, female fecundity via the induction of adventitious lateral root formation.

WOX11-mediated adventitious lateral root formation requires the transcription factor LATERAL ORGAN BOUNDARIES DOMAIN16 (LBD16). WOX11 directly binds to the promoter of LBD16, which induces the asymmetric radial expansion of root founder cells (Goh et al., 2012, Vilches Barro et al., 2019). Subsequently, founder cells divide asymmetrically to initiate an adventitious lateral root primordium. Here, we first investigated whether the transcriptional repressor mutant *35S:WOX11-SRDX* or the knockout *lbd16-2* mutant alter syncytium hypertrophy and female fecundity. Thus, we counted the number of nematodes that successfully established an infection, measured syncytium and female size, and quantified the number of eggs produced by females in the mutants compared to wild-type plants. Next, we conducted a correlation analysis to test for causality between WOX11-mediated adventitious lateral root formation and syncytium hypertrophy and female fecundity. Furthermore, we questioned whether WOX11 could directly affect the capacity of syncytia to accommodate large volumes of water and assimilates by modulating plant cell wall plasticity. To this end, we analyzed the effect of a chemical inhibitor of cellulose biosynthesis on syncytium hypertrophy in the *35S:WOX11-SRDX* and *lbd16-2* mutants compared to wild-type Arabidopsis. Finally, given that WOX11 has been previously implicated in the regulation of ROS (Liu *et al*., 2021, Wang *et al*., 2021, Xu *et al*., 2023) and that ROS are known to modulate cell wall plasticity (Eljebbawi *et al*., 2021), we researched whether WOX11 modulates ROS homeostasis in nematode syncytia. Our findings indicate that WOX11 attenuates female fecundity by restricting the hypertrophy of syncytial elements. Remarkably, this function of WOX11 in plant cell size control is not causally linked to adventitious lateral root formation. Instead, WOX11 may modulate syncytium hypertrophy via ROS-mediated cell wall plasticity mechanisms.

## Materials and Methods

### Plant material and growth conditions

The Arabidopsis (*Arabidopsis thaliana*) lines Col-0, *35S::WOX11-SRDX* (Liu *et al.,* 2014), and *lbd16-2* (Fan *et al.,* 2012) were used. The transcriptional repressor *35S::WOX11-SRDX* mutant was chosen since it was reported to be more strongly impaired in adventitious lateral root formation compared to the double mutant *wox11-2 wox12-1* (Liu *et al*., 2014, Sheng *et al*., 2017). For *in vitro* experiments, Arabidopsis seeds were vapor sterilized and grown on modified Knop medium (Sijmons *et al*., 1991) in a growth chamber with a 16 h : 8 h, light : dark photoperiod at 21°C. Plants were grown horizontally in 12-well plates for the *in vitro* infection assay and vertically in 12×12 cm Petri dishes for microscopy experiments. For pot experiments, non-sterile Arabidopsis seeds were sown on top of silver sand and grown under greenhouse conditions with 19-21 ⁰C and a 16 h : 8 h, light : dark photoperiod.

### Nematode sterilization

*Heterodera schachtii* (Woensdrecht population from IRS, The Netherlands) cysts were obtained from infected *Brassica oleracea* roots grown in sand (Baum *et al*., 2000). The cysts were hatched for seven days in a solution containing 1.5 mg ml^-1^ gentamycin sulfate, 0.05 mg ml^-1^ nystatin, and 3 mM ZnCl_2_. Next, *H. schachtii* second-stage juveniles (J2s) were separated from debris using a 35% sucrose gradient and incubated in a sterilization solution (0.16 mM HgCl_2_, 0.49 mM NaN_3_, and 0.002% Triton X-100) for 15 minutes. Finally, the J2s were washed three times with sterile tap water and re-suspended in 0.7% Gelrite (Duchefa Biochemie, Haarlem, The Netherlands).

### Pot experiment

21-day-old Arabidopsis plants were inoculated with 25 non-sterile J2s per gram of dry sand. This inoculation density was previously found to yield enough cysts (∼10) without causing excessive inhibition of plant growth (Fig. **S1**). Distribution of the pots containing the different genotypes in the trays followed a randomized block design. At 28 days post inoculation (dpi), watering of the plants was discontinued and sand within the pots was allowed to dry for one month. The shoots were cut off and each pot was wrapped in aluminum foil and autoclaved. Samples were sent to the NAK (Emmeloord, The Netherlands) for automated cyst extraction. Cysts were crushed and the number of eggs and J2s per cyst was counted following the protocol by Teklu *et al*. (2018).

### Isoxaben treatment

The procedure for the isoxaben treatment was adapted from literature (Chaudhary *et al*., 2020). Four-day-old Arabidopsis seedlings were inoculated with 15 J2s per seedling or with a mock solution. At 5 dpi, seedlings were transferred to 55 mm round petri dishes containing 10 ml of liquid Knop medium and either 600 nM isoxaben or 0.01% DMSO. After 5 hours of treatment, seedlings were transferred to fresh liquid Knop medium. Nematode syncytia were imaged by brightfield microscopy as described in the next section. Roots were mounted in 10 μg ml^−1^ propidium iodide staining to image the elongation zone of non-infected seedlings and pictures were taken using a Leica SP8 confocal microscope (excitation/emission 488/600-640 nm). For each sample, the one dimension size (maximum width) of five epidermal cells at the elongation zone was measured using Fiji software (Schindelin *et al*., 2012), after which the average width was calculated.

### Measurement of female and syncytium size

Pictures of mature females and syncytia were taken at 28 dpi using an Olympus SZX10 binocular (Olympus, Tokyo, Japan) with a ×1.5 objective and ×2.5 magnification. For the observation of syncytia at 5 dpi, seedlings were mounted in water and imaged with an Axio Imager.M2 light microscope (Zeiss) via a ×20 objective and a differential interference contrast filter. Images were taken with an AxioCam MRc5 camera (Zeiss). The size of females and syncytia was extracted from the pictures by manually measuring the maximum two-dimensional surface areas using Fiji software (Schindelin *et al*., 2012). The length of syncytia was taken by drawing a longitudinal line in the middle of the syncytium. The width was measured by selecting the widest point of the syncytium.

### DAB staining

Five days after inoculation, infected and non-infected Arabidopsis seedlings were stained with DAB as described previously (Siddique *et al*., 2014). First, seedlings were incubated in a DAB staining solution (10 mg ml^-1^ in water) for 2 hours in the dark. Then, they were bleached using an ethanol: lactic acid: glycerol (3:1:1) solution. Finally, seedlings were mounted in water and imaged with an Axio Imager.M2 light microscope (Zeiss) via a ×20 objective and a differential interference contrast filter. DAB staining intensity was scored on a scale from zero to six as described in Fig. **S2**.

### Statistical analyses

Data was analysed using the R software version 3.6.3 (Windows, x64). The R packages used were tidyverse (https://CRAN.R-project.org/package=tidyverse), ARTool (https://CRAN.R-project.org/package=ARTool) and multcompView (https://CRAN.R-project.org/package=multcompView). Correlation between variables was calculated using Spearman Rank-Order Correlation coefficient. The 95% confidence interval of the linear models was calculated using geom_smooth in R. For normally distributed data, significance of the differences among means was calculated by ANOVA followed by Tukey’s HSD test for multiple comparisons. For non-parametric two-factorial ANOVA, an Aligned Rank Transform followed by an ART-contrast test for multiple comparisons was performed.

## Results

### WOX11 restricts the radial expansion of nematode syncytia and attenuates female growth

In a previous study, we showed that WOX11 modulates plant tolerance to infections by endoparasitic cyst nematodes (Willig *et al*., 2023). Here, we asked whether this also affects nematode parasitism. We first tested whether WOX11 and its downstream target LBD16 affect the number of nematodes that successfully establish an infection. Therefore, we inoculated 14-day-old Col-0, *35S::WOX11-SRDX* (transcriptional repressor), and *lbd16-2* plants with the beet cyst nematode *Heterodera schachtii*. At 28 days post inoculation (dpi), we counted the number of females and males per plant. None of the mutants differed from wild-type Col-0 in the number of females or males (Fig. **1a, b**). Next, we investigated whether WOX11 and LBD16 affect syncytium expansion and female growth. For this purpose, pictures of nematode syncytia with only one female were taken from infected Col-0, *35S::WOX11-SRDX*, and *lbd16-2* plants at 28 dpi. The maximum two-dimensional surface area of the adult females and their corresponding syncytia was measured in the focal plane. This showed that the mutant plants *35S::WOX11-SRDX* and *lbd16-2* had significantly larger females and syncytia than wild-type Col-0 (Fig. **1c-e**). To further understand in which direction (longitudinal or radial) syncytium expansion in the mutants differed from Col-0, we also measured the width and length maxima of syncytia. We found that *35S::WOX11-SRDX* and *lbd16-2* had significantly wider syncytia compared to wild-type plants (Fig. **1f**). In contrast, no difference in syncytium length was observed among the genotypes (Fig. **1g**). We concluded that WOX11 and LBD16 restrict the radial expansion of syncytial elements and attenuate female growth.

**Fig. 1.**
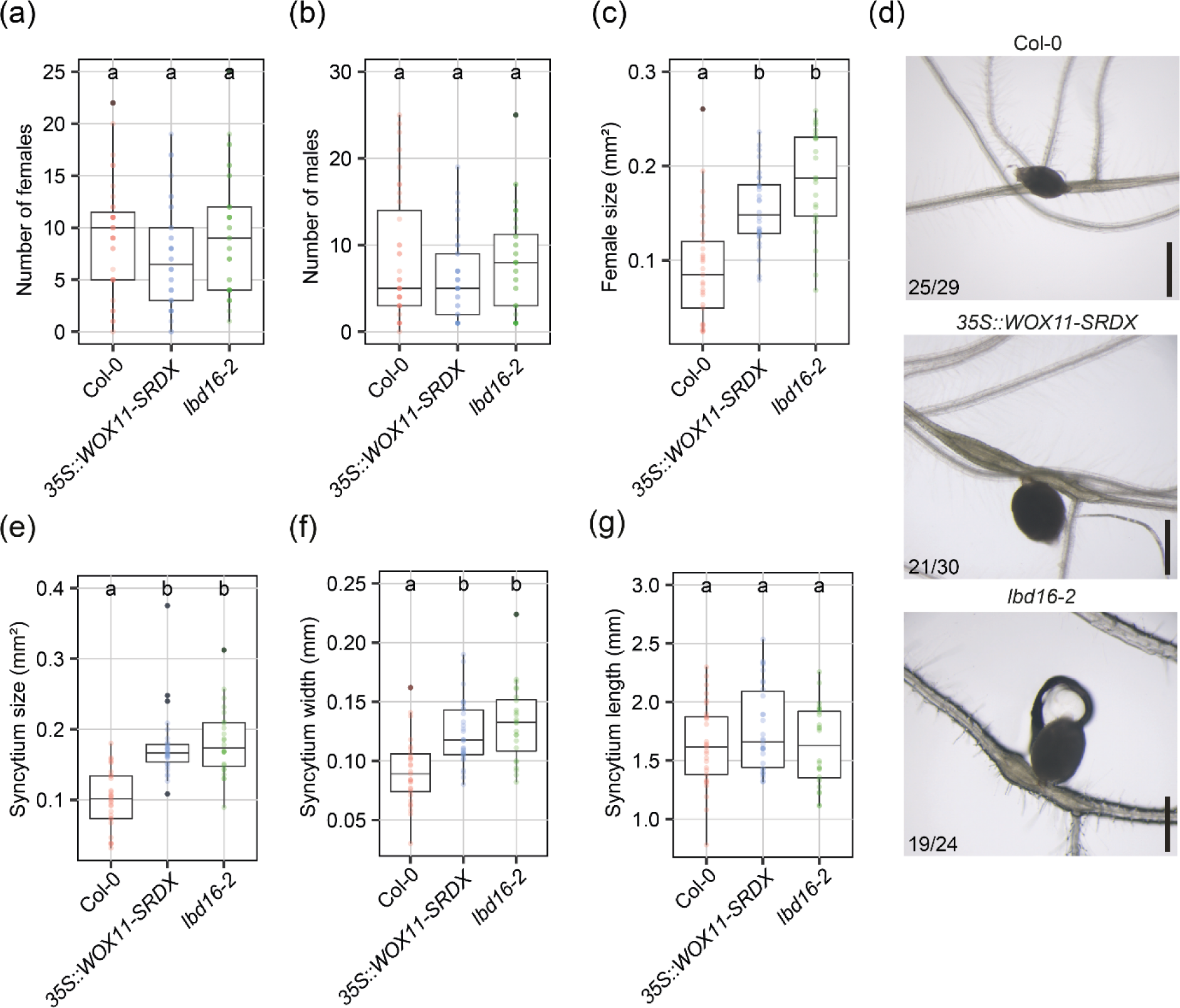
WOX11 and LBD16 restrict *Heterodera schachtii* syncytium hypertrophy and attenuate female growth. 14-day-old Col-0, *35S::WOX11-SRDX*, and *lbd16-2* Arabidopsis plants were inoculated with 250 J2s per plant. **(a)** The number of females and **(b)** the number of males were counted at 28 dpi (n=36). **(c)** Female size, **(e)** syncytium size, and **(f)** syncytium width, and **(g)** length were measured at 28 dpi (n=24-30). This experiment was performed three times with similar outcomes and data was pooled for statistical analysis. Significance of differences between genotypes was calculated by ANOVA followed by Tukey’s HSD test for multiple comparisons. Different letters indicate statistically different groups (*P*<0.05). **(d)** Representative pictures of females and syncytia at 28 dpi. Numbers at the bottom left corner indicate how often a similar phenotype as shown in the representative pictures was observed. Scale bar is 0.5 mm.

### WOX11 attenuates nematode female fecundity in soil

Nematode assays *in vitro* can yield different results from experiments in soil, where plant growth is subjected to additional stresses and more variable conditions (Grenier *et al*., 2020). To assess if WOX11 and LBD16 may influence female growth and thus fecundity in soil, we cultivated *35S::WOX11-*SRDX, *lbd16-2,* and wild-type Arabidopsis plants in pots with silver sand and then inoculated them with *H. schachtii.* At 28 dpi, the sand in the pots was left to dry, after which the dead females, referred to as cysts, were extracted from the sand, counted, and crushed to count the number of eggs or J2s within each cyst (Fig. **2**). The number of cysts from *35S::WOX11-SRDX* plants was not significantly different from the wild-type Col-0. However, *lbd16-2* plants had a lower number of cysts per plant compared to Col-0, suggesting that LBD16 may enhance plant susceptibility to nematode infections in pot conditions (Fig. **2a**). Interestingly, we found that the average number of eggs or J2s per cyst was significantly higher in *35S::WOX11-SRDX* compared to Col-0 plant. Although *lbd16-2* also showed a higher average number of eggs or J2s per cyst than Col-0, this difference was not significant (Fig. **2b**). Thus, we concluded that WOX11 does not affect plant susceptibility to nematode infection but restricts nematode female fecundity in soil.

**Fig. 2.**
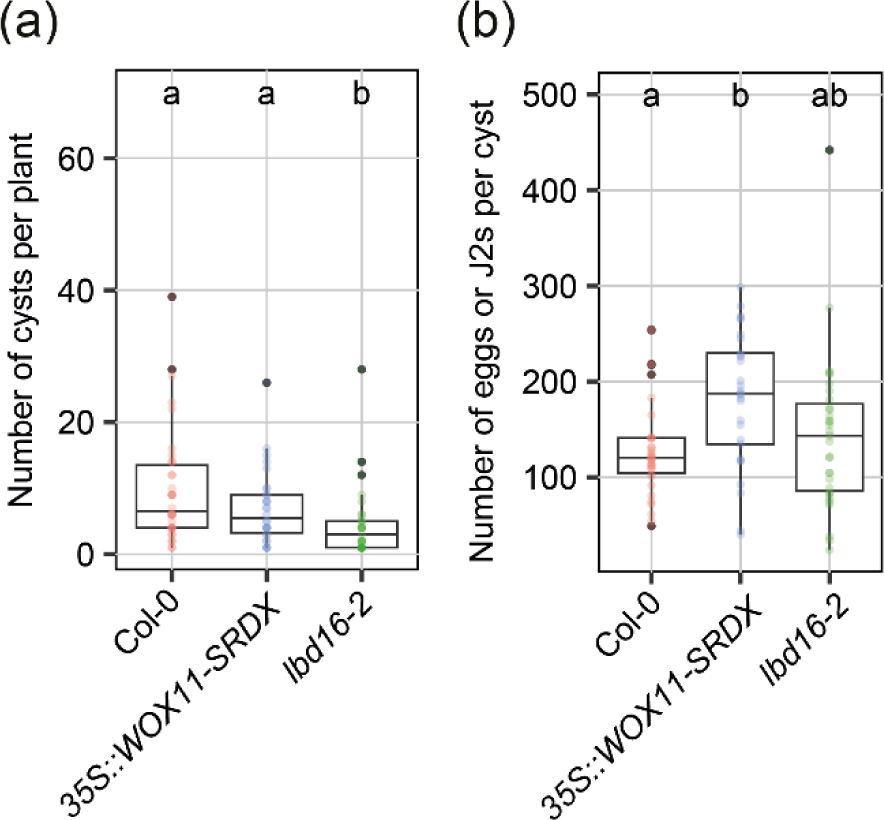
WOX11 attenuates *Heterodera schachtii* female fecundity in soil. 21 days after growing wild-type Col-0 and *35S::WOX11-SRDX* and *lbd16-2* mutants on silver sand, plants were inoculated with 25 J2s/g of dry sand. Pot disposition in the trays followed a randomized block design. At 28 dpi, cysts were extracted, counted, and crushed to count the number of eggs or J2s contained in each cyst. **(a)** Number of cysts per plant. **(b)** Average number of eggs or J2s per cyst. Significance of differences between genotypes was calculated by ANOVA followed by Tukey’s HSD test for multiple comparisons. Different letters indicate statistically different groups (*P*<0.05, n=30-35).

### The effect of WOX11 on syncytium hypertrophy and female fecundity is not causally linked to adventitious lateral root formation

WOX11-adventitious lateral roots emerge in proximity or adjacent to nematode syncytia (Golinowski *et al*., 1996, Willig *et al*., 2023). As adventitious lateral roots are both strong sinks of assimilates (Stitz *et al*., 2023) and an additional source of water and minerals (Levin *et al*., 2020), we hypothesized that they could either compete or support syncytium hypertrophy and nematode fecundity. To test this, we counted the number of adventitious lateral roots emerging from 28 dpi syncytia in Col-0, *35S::WOX11-*SRDX, and *lbd16-2* and performed a correlation analysis with female and syncytium size. Interestingly, we found a positive correlation between the number of adventitious lateral roots emerging from syncytia and both syncytium and female size in wild-type Col-0 (Fig. **3a, b**). Thus, the emergence of adventitious lateral roots may support syncytium hypertrophy and female fecundity. However, this positive correlation was not observed in the *35S::WOX11-*SRDX and *lbd16-2* mutants (Fig. **3a, b**). This indicates that adventitious lateral root formation at nematode syncytia is not causally linked to the role of WOX11 in syncytium hypertrophy and female fecundity.

**Fig. 3.**
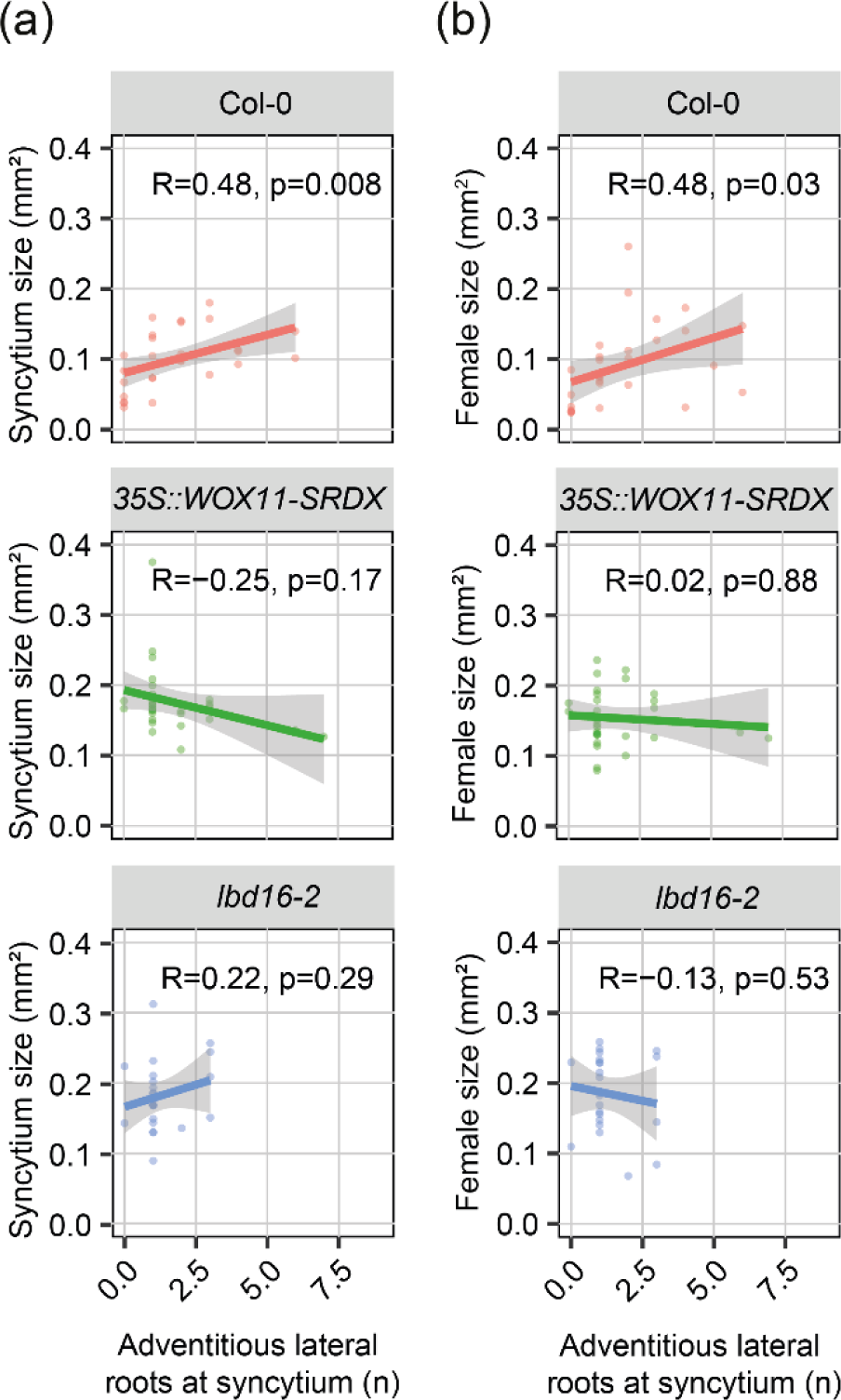
The effect of WOX11 and LBD16 on *Heterodera schachtii* syncytium hypertrophy and female fecundity is not causally linked to adventitious lateral root formation. 14-day-old Col-0, *35S::WOX11-SRDX*, and *lbd16-2* plants were inoculated with 250 *H. schachtii* J2s per plant. At 28 dpi, the number of adventitious lateral roots in contact with nematode syncytia was counted and the size of females and syncytia was measured. **(a)** Correlation between the number of adventitious lateral roots and syncytium size. **(b)** Correlation between the number of adventitious lateral roots and female size. This experiment was performed three times with similar outcomes and data was pooled for statistical analysis. Correlation (R) between two variables was calculated using Spearman’s rank-order correlation coefficient (n=24-30). Gray area indicates 95% confidence interval.

### WOX11 restricts the early radial expansion of syncytial elements

Radial expansion of nematode syncytia is mainly determined by the cellular hypertrophy of syncytial elements at 5 dpi (Magnusson and Golinowski, 1991, Golinowski *et al*., 1996). At this stage, syncytial elements are symplastically isolated from the surrounding host tissue, which likely enables the build op of turgor pressure required for cellular hypertrophy (Ruan *et al*., 2004, Hofmann *et al*., 2007). Since we previously observed that WOX11 and LBD16 are expressed between 2 and 7 dpi (Willig *et al*., 2023), we asked whether WOX11 and LBD16 play a role in the early phases of syncytium expansion. To this end, we measured syncytium width in *35S::WOX11-SRDX* and *lbd16-2* mutants at 5 dpi. We found that, already at such an early time point, the two mutants displayed wider syncytia compared to the wild-type Col-0 (Fig. **4a, b**). Thus, WOX11 and LBD16 limit the early radial expansion of syncytial elements.

**Fig. 4.**
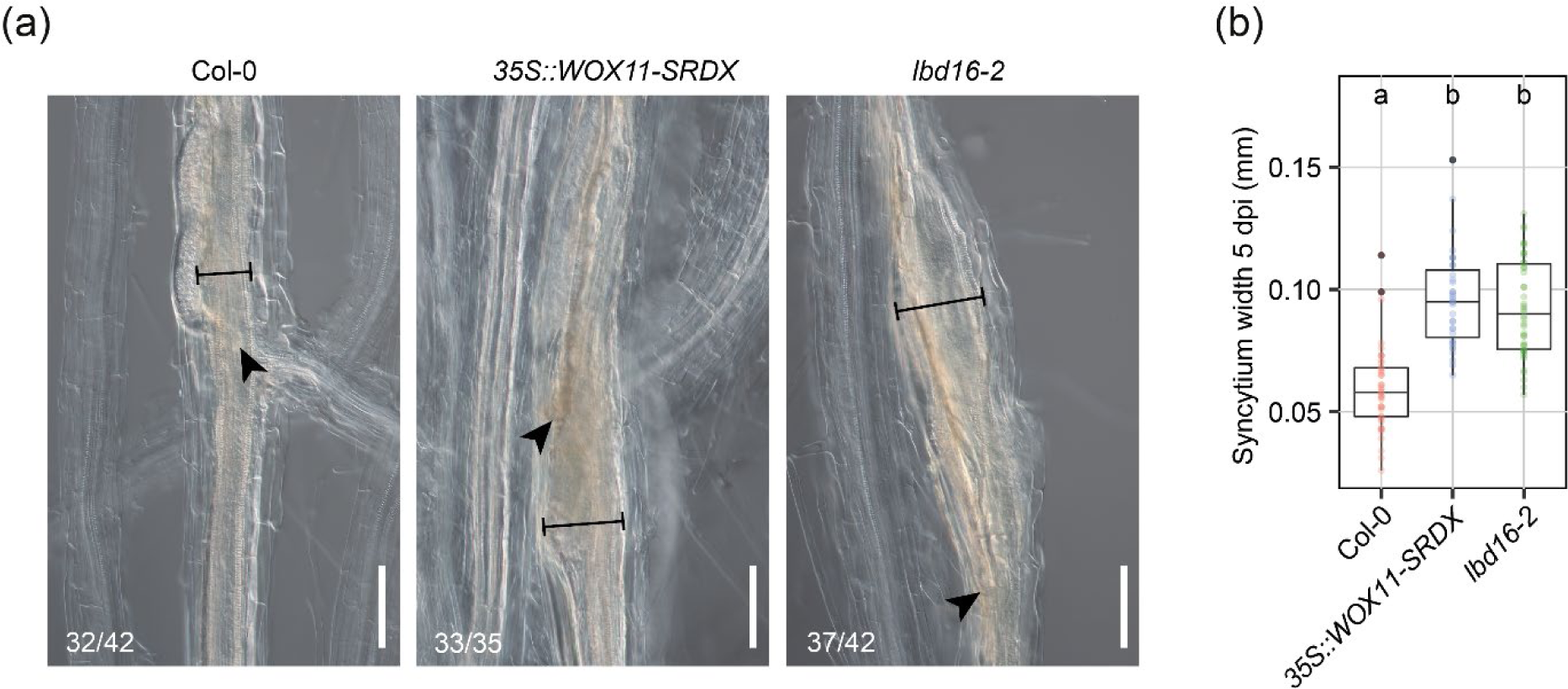
WOX11 restricts the early radial expansion of *Heterodera schachtii* syncytial elements. Four-day-old Arabidopsis seedlings were inoculated with 15 *H. schachtii* J2s per plant. At 5 dpi, pictures were taken of syncytia and width was measured. **(a)** Representative pictures of syncytia at 5 dpi. Black arrowheads indicate the nematode head. Black lines indicate syncytium width. Numbers at the bottom left corner indicate how often a similar phenotype as shown in the representative pictures was observed. Scale bar is 100 μm. **(b)** Syncytium width at 5 dpi. This experiment was performed three times with similar outcomes and data was pooled for statistical analysis. Significance of differences between genotypes was calculated by ANOVA followed by Tukey’s HSD test for multiple comparisons. Different letters indicate statistically different groups (*P*<0.001, n=35-42).

### WOX11 restricts the cell wall extensibility of syncytial elements

Root cells in the maturation zone are typically long and narrow. This is because root cells elongate longitudinally in the root elongation zone before completing differentiation and entering the maturation zone (Verbelen *et al*., 2006). The orientation of cellulose microfibrils and the properties of other cell wall components increase the stiffness of the lateral cell walls, thus guiding the expansion in the longitudinal direction (Chaudhary *et al*., 2020). However, plant developmental processes and environmental stresses can cause changes in cell wall structure and composition, leading to enhanced radial expansion of root cells (Gigli-Bisceglia *et al*., 2020). It was previously shown that LBD16 regulates the asymmetric radial expansion of founder cells, which involves the differential organization of cortical microtubules driving the deposition of cellulose microfibrils (Vilches Barro *et al*., 2019). Here, we hypothesized that WOX11 and LBD16 restrict the radial expansion of syncytial elements by modulating cell wall plasticity mechanisms.

To test this, we inhibited cellulose biosynthesis using isoxaben, a chemical that causes the internalization of cellulose synthase complexes from the plasma membrane to cytosolic vesicles (Tateno *et al*., 2016). Disruption of cellulose biosynthesis or changes in cellulose microfibril alignment affects the directional growth of plant cells, which causes the cells to become radially swollen. Hence, four-days-old Arabidopsis *35S::WOX11-SRDX, lbd16-2,* and wild-type Col-0 seedlings were inoculated with *H. schachtii.* At 5 dpi, seedlings were treated either with isoxaben or DMSO as a negative control. Isoxaben treatment led to a significant increase in syncytium width in wild-type Col-0 plants, which phenocopied the DMSO-treated syncytia in the *35S::WOX11-SRDX* and *lbd16-2* mutants (Fig. **5a, b**). Thus, inhibition of cellulose biosynthesis causes similar effects on radial expansion of syncytial elements as a disruption in WOX11- and LBD16-mediated pathways.

**Fig. 5.**
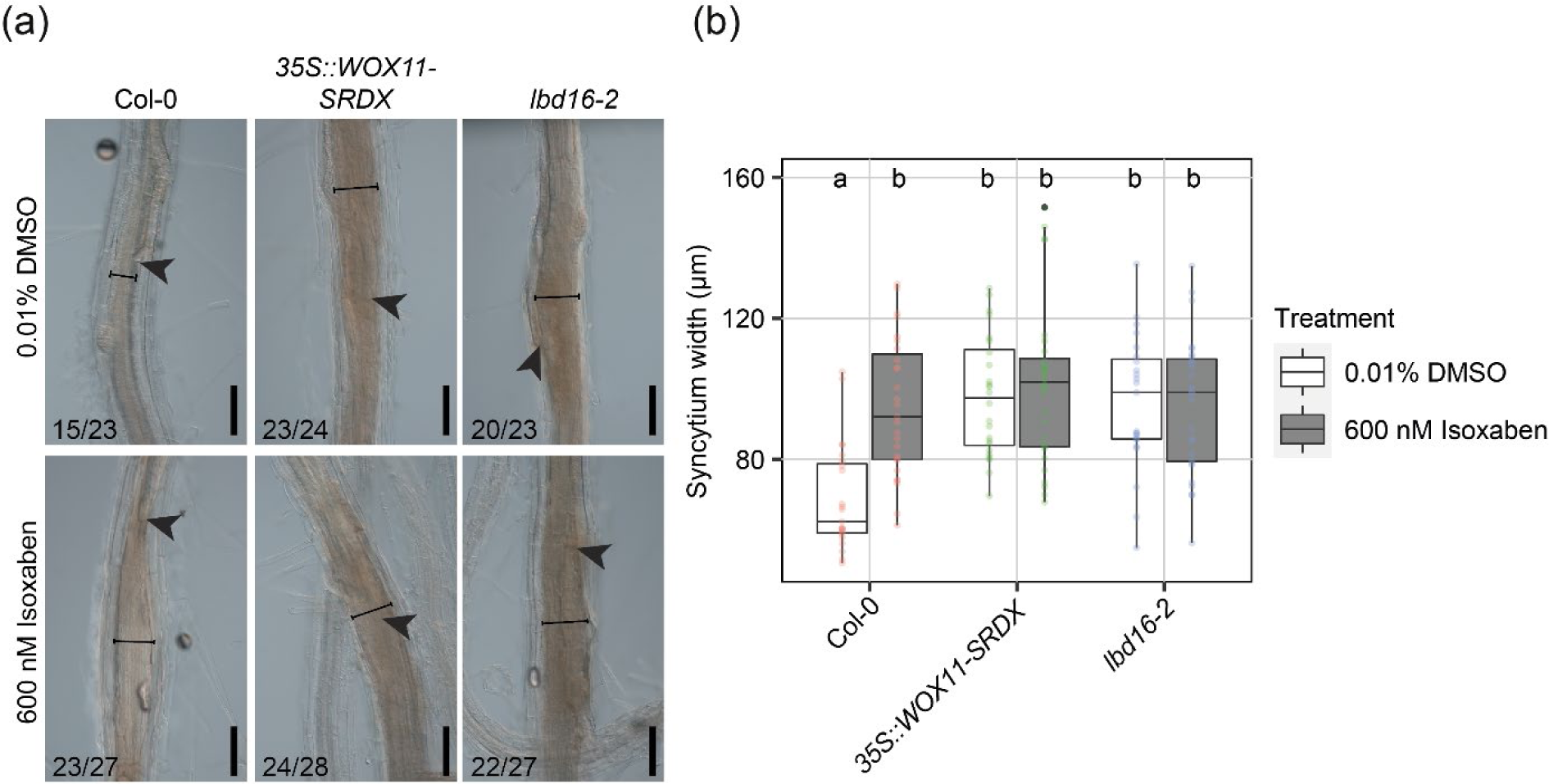
Chemical inhibition of cellulose biosynthesis in Col-0 phenocopies the enhanced radial expansion of syncytial elements observed in *35S::WOX11-SRDX* and *lbd16-2*. Four-day-old Col-0, *35S::WOX11-SRDX,* and *lbd16-2* Arabidopsis seedlings were inoculated with 15 *H. schachtii* J2s. At 5 dpi, seedlings were transferred to liquid Knop medium containing either 600 nM isoxaben (ISX) or 0.01% DMSO. **(a)** Representative pictures of nematode syncytia. Black arrowheads indicate the nematode head. Black lines indicate syncytium width. Numbers at the bottom left corner indicate how often a similar phenotype as shown in the representative pictures was observed. Scale bar is 100 μm. **(b)** Quantification of syncytium width. This experiment was performed two times with similar outcomes and data was pooled for statistical analysis. Significance of differences between genotypes was calculated by ANOVA followed by Tukey’s HSD test for multiple comparisons (*P*<0.05, n=23-27). Different letters indicate statistically different groups.

Interestingly, isoxaben treatment did not have a visible additive effect on the radial expansion of syncytia in *35S::WOX11-SRDX* and *lbd16-2* (Fig. **5a, b**), suggesting that the syncytial cell walls in the mutants already reached their maximum extensibility. Therefore, we hypothesized that WOX11 and LBD16 attenuate plant cell wall extensibility. We verified this by treating nine-day-old non-infected seedlings with isoxaben, which is known to cause the radial expansion of root epidermal cells at the elongation zone (Chaudhary *et al*., 2020). After 5 hours of treatment with isoxaben, we found that the width of epidermal cells at the elongation zone of *35S::WOX11-SRDX* and *lbd16-2* increased dramatically compared to wild-type Col-0 (Fig. **6a, b**). Notably, the width of epidermal cells in all genotypes was similar upon treatment with the negative control (Fig. **6a, b**). Altogether, our data suggests that WOX11 and LBD16 restrict cell wall extensibility of both isoxaben-treated epidermal cell at the elongation zone and syncytial elements in the mature zone.

**Fig. 6.**
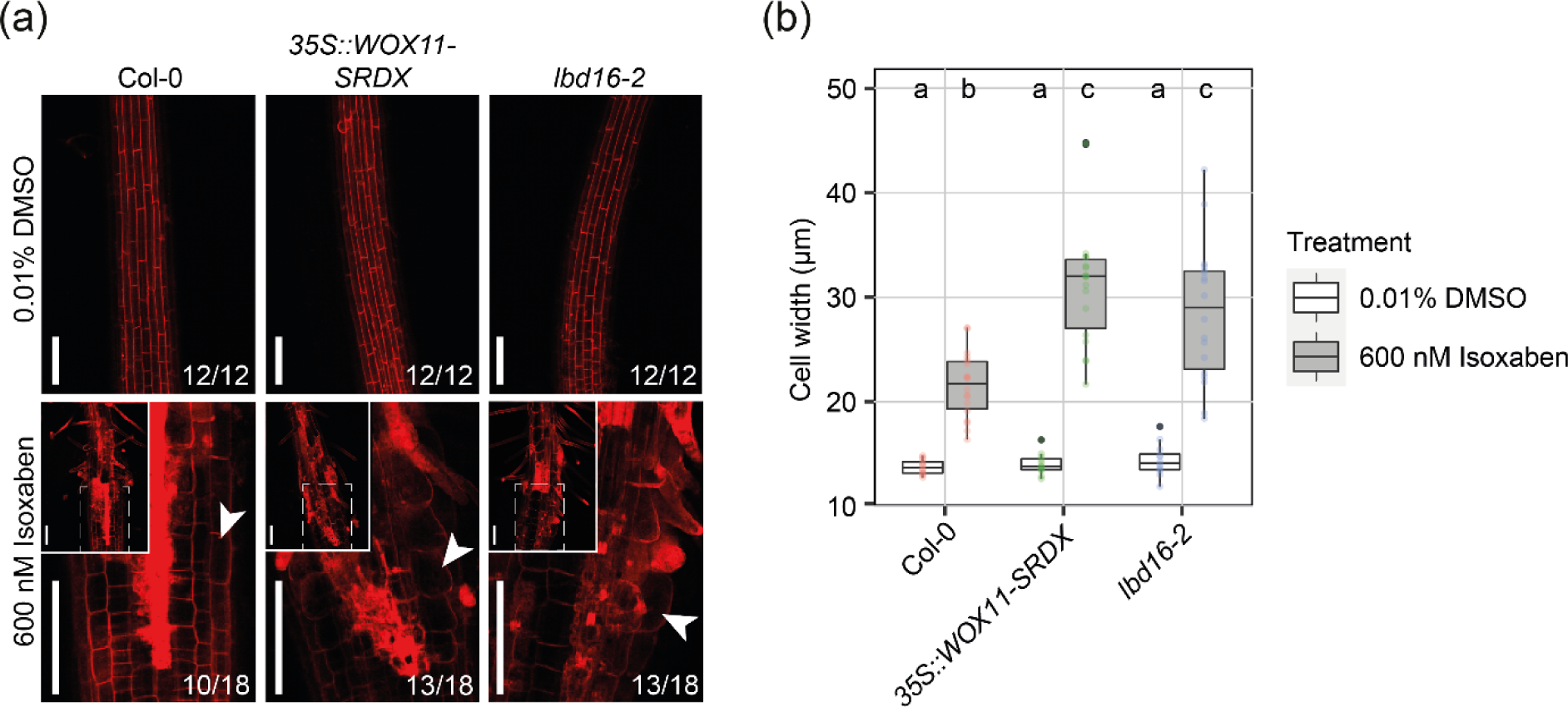
WOX11 restricts the radial expansion of epidermal cells at the elongation zone in response to isoxaben treatment. Nine-day-old Col-0, *35S::WOX11-SRDX,* and *lbd16-2* Arabidopsis seedlings were transferred to liquid Knop medium containing either 600 nM isoxaben or 0.01% DMSO. After 5 hours, seedlings were mounted in 10 μg ml−1 propidium iodide for imaging. **(a)** Representative pictures of the elongation zone. White arrowheads indicate a radially expanded epidermal cell. The top-left inserts represent the original zoomed-out pictures. Numbers at the bottom left corner indicate how often a similar phenotype as shown in the representative pictures was observed. Scale bar is 100 μm. **(b)** Quantification of epidermal cells width. Each data point corresponds to the average width of five epidermal cells for each root sample. This experiment was performed two times with similar outcomes and data was pooled for statistical analysis. Significance of differences between genotypes was calculated by ANOVA followed by Tukey’s HSD test for multiple comparisons (*P*<0.0001, n=12-18). Different letters indicate statistically different groups.

### WOX11 modulates ROS homeostasis in nematode syncytia

ROS homeostasis is important in determining plant cell wall plasticity (Eljebbawi *et al*., 2021). WOX11 was previously found to regulate ROS homeostasis for crown root development in rice (Xu *et al*., 2023) and during drought and salt stress in poplar (Liu *et al*., 2021, Wang *et al*., 2021). Therefore, we hypothesized that WOX11-mediated plant cell wall plasticity involves the regulation of ROS. First, we stained 5 dpi syncytia in the *35S::WOX11-SRDX* and *lbd16-2* mutants with a DAB solution, which is oxidized by peroxidases in the presence of H_2_O_2_ (Eljebbawi *et al*., 2021). This revealed that nematode syncytia in the *35S::WOX11-SRDX* and *lbd16-2* mutants have significantly lower ROS compared to wild-type Col-0 (Fig. **7a, b**). Thus, we concluded that WOX11 modulates ROS homeostasis at nematode syncytia.

**Fig. 7.**
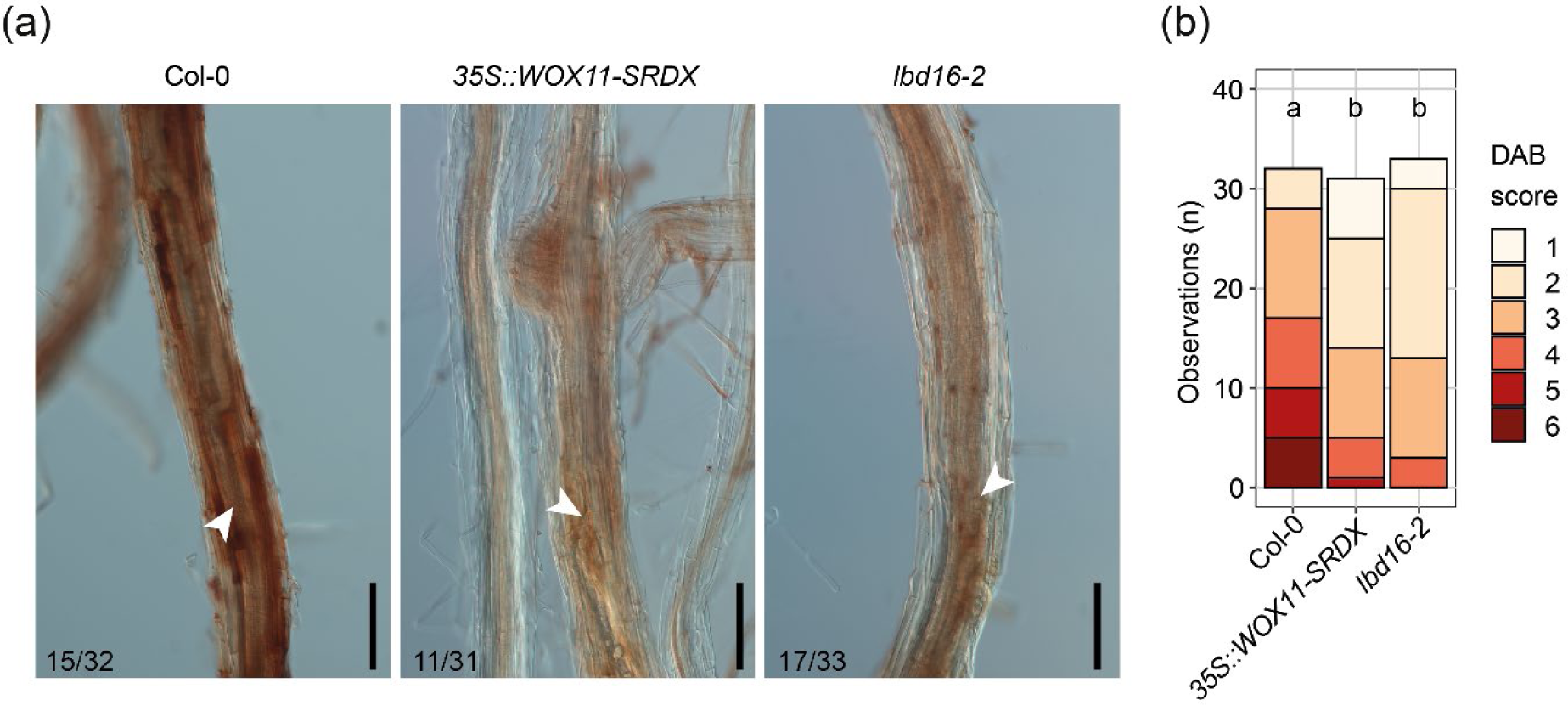
WOX11 and LBD16 increase ROS levels in *Heterodera schachtii* syncytia. Four-day-old Col-0, *35S::WOX11-SRDX* and *lbd16-2* Arabidopsis seedlings were inoculated with 15 *H. schachtii* J2s. At 5 dpi, seedlings were incubated in a DAB staining solution for 2 hours in the dark. **(a)** Representative pictures of nematode syncytia. White arrowheads indicate the nematode head. Numbers at the bottom left corner indicate how often a similar phenotype as shown in the representative pictures was observed. Scale bar is 100 μm. **(b)** Scoring of DAB staining intensity on a scale from 1 to 6, based on the pictures shown in Figure S2. Bar graphs indicate how often a certain DAB score is observed in each genotype. This experiment was performed three times with similar outcomes and data was pooled for statistical analysis. Significance of differences between genotypes was calculated by and Aligned Ranks Transform (ART) non-parametric ANOVA followed by an ART-contrast test for multiple comparisons (*P*<0.0001, n=31-35). Different letters indicate statistically different groups.

**Fig. 8.**
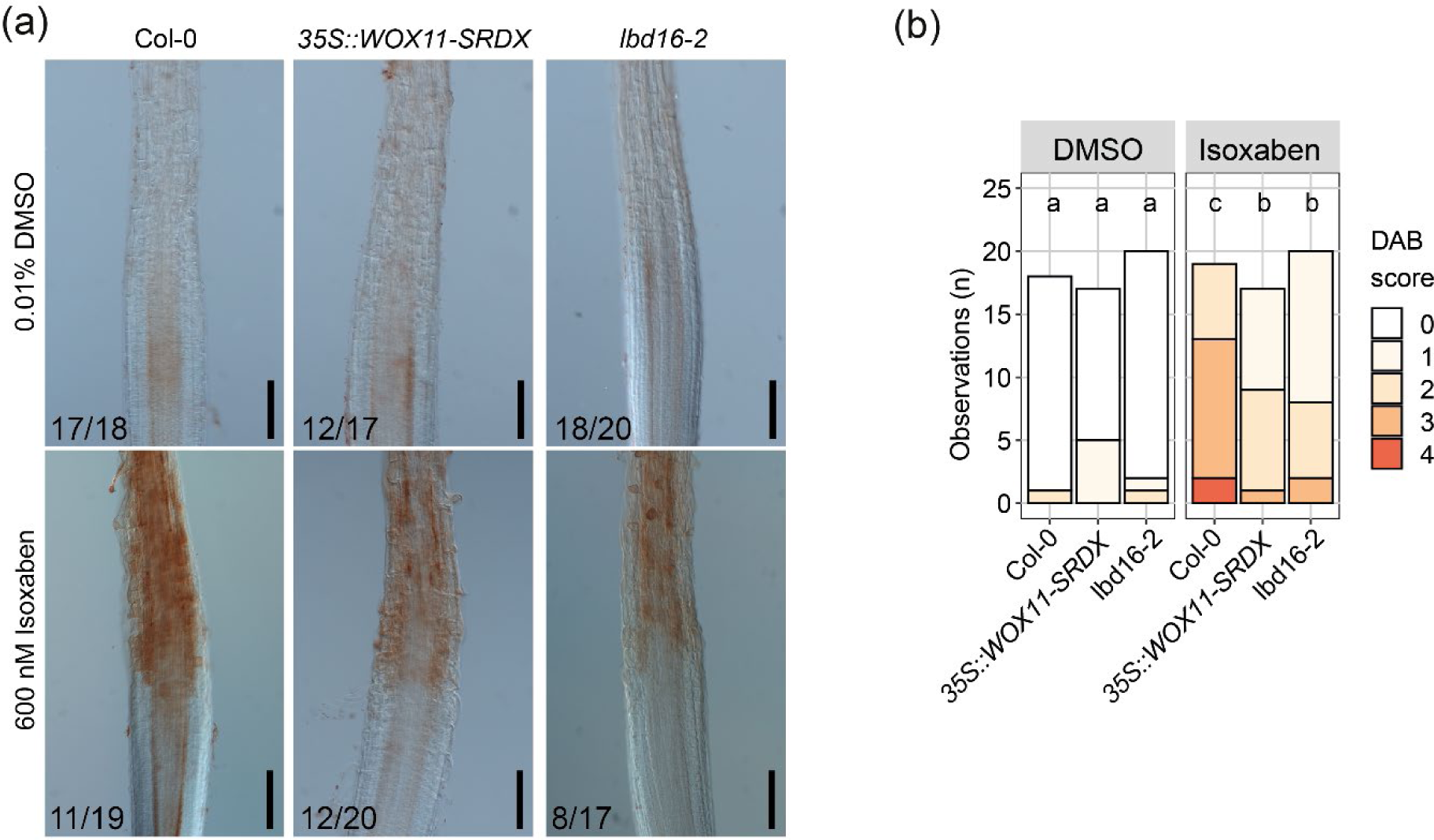
WOX11 and LBD16 increase ROS levels at the elongation zone in response to cellulose biosynthesis inhibition. 9-day-old Col-0, *35S::WOX11-SRDX* and *lbd16-2* Arabidopsis seedlings were transferred to liquid KNOP medium containing either 600 nM isoxaben or 0.01% DMSO. After 5 hours, seedlings were incubated in a DAB staining solution for 2 hours in the dark. **(a)** Representative pictures of the elongation zone. Numbers at the bottom left corner indicate how often a similar phenotype as shown in the representative pictures was observed. Scale bar is 100 μm. **(b)** Scoring of DAB staining intensity on a scale from 0 to 4, based on the pictures shown in Figure S2. Bar graphs indicate how often a certain DAB score is observed in each genotype. This experiment was performed two times with similar outcomes and data was pooled for statistical analysis. Significance of differences between genotypes was calculated by and Aligned Ranks Transform (ART) non-parametric ANOVA followed by an ART-contrast test for multiple comparisons (*P*<0.0001, n=18-20). Different letters indicate statistically different groups.

To verify whether WOX11-mediated ROS burst is linked to cell wall plasticity responses, we treated non-infected nine-day-old Col-0, *35S::WOX11-SRDX*, and *lbd16-2* seedlings with 600 nM isoxaben or 0.01% DMSO for 5 hours, followed by DAB staining. Isoxaben treatment caused an increase in ROS at the elongation zone in all genotypes. However, the levels of ROS in the *35S::WOX11-SRDX* and *lbd16-2* mutants were significantly lower compared to Col-0 (Fig. **6a, b**). Thus, we concluded that modulation of ROS homeostasis is a plausible mechanism underlying WOX11-mediated cell wall plasticity at nematode syncytia.

## Discussion

Plant developmental plasticity mitigates the negative impacts of cyst nematode infections on growth, yet its potential impact on nematode parasitism remains largely unknown. We recently reported that plant perception of cyst nematode invasion induces the formation of WOX11-adventitious lateral roots at nematode infection sites (Guarneri *et al*., 2023, Willig *et al*., 2023). Through this local root plasticity response, WOX11 compensates for the inhibition of primary root growth caused by nematode infection, which benefits overall plant growth and development (Willig *et al*., 2023). In this study, we shift perspective from the plant to the nematode and investigate whether WOX11-mediated developmental plasticity has an impact on nematode fecundity. Our findings support a model where WOX11 modulates ROS homeostasis and cell wall plasticity mechanisms that attenuate syncytial cell size and nematode fecundity.

We provide evidence that WOX11 restricts cyst nematode female growth and egg production. Although it was previously found that WOX11 decreases host invasion by infective juveniles (Willig *et al*., 2023), the repressor *35S::WOX11-SRDX* did not alter the number of nematodes successfully establishing an infection. Yet, Arabidopsis roots of the *35S::WOX11-SRDX* repressor displayed bigger females compared to wild-type Col-0 *in vitro.* When plants cultivated in soil were inoculated with *H. schachtii*, the cysts extracted from *35S::WOX11-SRDX* contained a higher number of eggs and J2s as compared to the cysts obtained from wildtype Col-0.

Consistently, the downstream target of WOX11, LBD16, similarly decreased female fecundity. *In vitro, lbd16-2* showed the same trend as the repressor *35S::WOX11-SRDX,* displaying bigger females compared to wild-type Col-0. Additionally, LBD16 had no effect on the number of nematodes that successfully established an infection. This is in agreement with a previous study, where the repressor mutant *35S::LBD16-SRDX* did not alter the number of nematode-induced syncytia after inoculation with *H. schachtii in vitro* (Cabrera *et al*., 2014). In soil, *lbd16-2* showed a lower number of eggs or J2s per cyst compared to Col-0, albeit that this difference was not statistically significant. However, the number of cysts extracted from *lbd16-2* plants was lower compared to Col-0 and *35S::WOX11-SRDX*. LBD16 does not only regulate adventitious rooting but also lateral root formation (Sheng *et al*., 2017). Thus, the disruption of both rooting pathways in the *lbd16-2* may have decreased the chances of nematodes locating or penetrating the roots in our pot experiment. Moreover, the outcomes of nematode bioassays in soil are generally more variable and have a lower resolution than *in vitro* assays. Thus, the overall low infection rates of *lbd16-2* plants may underly the absence of statistical significance in the difference between the average number of eggs or J2s per cysts in *lbd16-2* and Col-0.

We found that WOX11-induced adventitious lateral root formation is not causally linked to WOX11-mediated syncytium hypertrophy and nematode fecundity. Our data showed that adventitious lateral roots nonetheless support syncytium hypertrophy and female growth. This is in line with a previous hypothesis that adventitious lateral roots may increase water influx towards nematode syncytia (Levin *et al*., 2020). However, the positive correlation between the number of adventitious lateral roots and syncytium and female size was disrupted in the repressor *35S::WOX11-SRDX* and in the *lbd16-2* mutants. This suggested that the role of WOX11 in modulating syncytium hypertrophy and female fecundity is separate from its role in adventitious lateral root formation.

WOX11 may attenuate female offspring size by directly restricting the availability of plant assimilates in nematode syncytia. We found that the repressor *35S::WOX11-SRDX* and *lbd16-2* enhanced the radial expansion of syncytia at 5 dpi, a stage where syncytial elements are symplastically isolated from the surrounding host tissue (Hofmann *et al*., 2007). During symplastic isolation, nematode syncytia take up sucrose from the phloem via the apoplast through active transporters, which attracts water from the xylem and builds up turgor pressure (Hofmann *et al*., 2007, Grundler and Hofmann, 2011). The cyclic feeding behavior of nematodes likely causes fluctuations of sucrose levels and turgor pressure that are accommodated by changes in cell wall plasticity and syncytium expansion (Zhang *et al*., 2017). If WOX11 restricts the radial expansion of syncytial elements, it could interfere with the ability of syncytia to accumulate assimilates and support the cyclic food demands of female nematodes.

Furthermore, our data suggests that WOX11 modulates the plasticity of syncytial cell walls. We found that inhibition of cellulose biosynthesis using isoxaben increased the radial expansion of syncytial elements similarly to the disruption of WOX11-mediated pathways. Moreover, in non-infected plants, WOX11 limited the radial expansion of epidermal cells at the elongation zone in response to isoxaben treatment. This indicates that WOX11 limits the extensibility of syncytial elements in the radial direction. Syncytial cell wall plasticity plays an important role in cyst nematode parasitism. Indeed, the silencing of two plant cell-wall modifying endo-B-1,4-glucanases reduced the number of females and their egg content in potato (Karczmarek *et al*., 2008). Endo-B-1,4-glucanases loosen the cell wall by modifying amorphous cellulose structures (Glass *et al*., 2015), which likely affects the expansion of syncytial elements. Moreover, the upregulation of many expansin proteins in nematode syncytia at 5 dpi suggests that syncytium expansion involves mechanisms of cell-wall loosening (Wieczorek *et al*., 2006). It is possible that WOX11 mediates cell wall plasticity in nematode syncytia by regulating the expression of expansin genes. Indeed, WOX11 was found to directly bind the promoter of an expansin gene and thereby modulate grain width in rice (Xiong *et al*., 2023).

Plant cell wall plasticity is a tightly regulated process involving many interacting components (Cosgrove, 2022). Organization and biosynthesis of cellulose microfibrils, the major load bearing components of the cell wall, are strongly influenced by cortical microtubules and by the properties and abundance of different polymers in the cell wall matrix (Li *et al*., 2015, Xiao *et al*., 2015, Du *et al*., 2020). Cortical microtubules guide cellulose synthase complexes, determining the organization of cellulose microfibrils (Li *et al*., 2015). In turn, cellulose deposition regulates microtubule organization through a positive feedback loop (Vilches Barro *et al*., 2019). Moreover, pectin methylation and hemicellulose were found to affect both cellulose biosynthesis and cortical microtubule organization (Xiao *et al*., 2015, Du *et al*., 2020). Due to this complex network, isoxaben treatment does not only cause internalization of cellulose synthase complex but also alters the organization of cortical microtubules (Vilches Barro *et al*., 2019). We observed that isoxaben treatment did not result in a measurable radial expansion of nematode syncytia in the *35S::WOX11-SRDX* and *lbd16-2* mutants. Interestingly, syncytium expansion is known to involve the disorganization of cortical microtubules (De Almeida Engler *et al*., 2004). Thus, one plausible explanation for our observation is that WOX11 and LBD16 mediate plant cell wall plasticity mechanisms that alter the organization of cortical microtubules. In this scenario, the level of microtubule organization in *35S::WOX11-SRDX* and *lbd16-2* syncytia would be too low to be further affected by isoxaben. Consistently, cortical microtubule organization was found to play an important role in LBD16-mediated asymmetric radial expansion of root founder cells (Vilches Barro *et al*., 2019).

Our study suggests that WOX11 modulates ROS homeostasis in nematode syncytia. We found that WOX11 and LBD16 increase ROS accumulation in 5 dpi nematode syncytia. This is in line with a recent study where WOX11 directly activates peroxidase activity to induce ROS production and regulate crown root development in rice (Xu *et al*., 2023). Besides, WOX11 was found to reduce cytotoxic levels of ROS in response to salt and drought stress in poplar (Liu *et al*., 2014, Wang *et al*., 2021). Thus, WOX11 modulates ROS homeostasis in different plant species and in response to multiple stresses. However, how WOX11-mediated ROS homeostasis affects plant developmental plasticity still remains unclear.

WOX11-mediated ROS homeostasis could modulate the extensibility of plant cell walls. ROS regulate both cell wall loosening and cell wall stiffening (Schmidt *et al*., 2016, Eljebbawi *et al*., 2021). H_2_O_2_ can be converted by peroxidases into OH^-^, which catalyzes the oxidative cleavage of hemicellulose and pectins in the apoplast and loosens the cell wall (Eljebbawi *et al*., 2021). Additionally, when H_2_O_2_ levels are high, peroxidases promote the oxidation and thereby cross-linking of extensins and phenolic compounds, leading to cell wall stiffness (Magliano and Casal, 1998, Brownleader *et al*., 2000, Eljebbawi *et al*., 2021). Furthermore, ROS homeostasis regulates tubulin polymerization during microtubule formation (Livanos *et al*., 2012), which could affect the orientation of cellulose biosynthesis and hereby change the cell wall structural properties (Li *et al*., 2015). We have observed that WOX11 induces ROS production at the elongation zone in response to cellulose biosynthesis inhibition by isoxaben. Moreover, WOX11 attenuates the radial expansion of epidermal cells at the elongation zone upon isoxaben treatment. Thus, we suggest that WOX11-mediated ROS homeostasis increases cell wall stiffness in response to cellulose biosynthesis inhibition. Similarly, WOX11 could modulate ROS to restrict syncytial cell wall extensibility. Whether this involves the regulation of cortical microtubules, the cross-linking of extensins and phenolics, or other cell wall plasticity mechanisms needs to be investigated.

We found that WOX11 increases ROS accumulation and attenuates female fecundity. However, previous research showed that the decreased ROS production in the *rboh*D/F mutant resulted in smaller syncytia and females compared to wild-type plants (Siddique *et al*., 2014; Chopra *et al*., 2021). While too high or too low levels of ROS can be deleterious for plants, modulation of ROS homeostasis within non-deleterious levels can regulate many cellular and physiological processes (Mittler, 2017, Willig *et al*., 2022). For instance, modulation of ROS homeostasis via WOX11 could possibly mediate changes in syncytial cell wall extensibility in response to the fluctuations in sucrose and turgor pressure due to nematode cyclic feeding behavior. This may regulate the volumes of water and assimilates available for nematode feeding and thereby affect female fecundity.

In conclusion, we showed that WOX11 controls cell size likely by modulating ROS homeostasis and cell wall plasticity. Furthermore, we demonstrated that as a result WOX11 attenuates female growth and offspring size. Our results point at a novel role of WOX11 in the plant response to nematode infection, which is distinct from WOX11-mediated adventitious rooting. Importantly, our findings provide evidence that plant developmental plasticity can modulate nematode parasitism.

## Acknowledgments

This work was supported by the Graduate School Experimental Plant Sciences (EPS). JJW is funded by Dutch Top Sector Horticulture & Starting Materials (TU18152). MGS was supported by NWO domain Applied and Engineering Sciences VENI grant (17282). JLLT was supported by NWO domain Applied and Engineering Sciences VENI (14250) and VIDI (18389) grants. No conflict of interest declared.

## Author contributions

GS, NG, and JJW conceived the project. NG and JJW designed the experiments and performed data collection. Data was analyzed and interpreted by NG, GS, JJW, JJLT, MGS, and PN. NG and GS wrote the article with inputs from AG, JJLT, JJW, MGS, PN, and VW.

## Data availability

The data that supports the findings of this study are available in the supplementary material of this article.

## Supporting information

Supporting Information Fig. S1 Number of *Heterodera schachtii* cysts at increasing inoculation densities in wild-type Col-0 grown in pots.

Supporting Information Fig. S2 Scale used for scoring the DAB staining of Arabidopsis roots.

Supporting Information Fig. S3 DAB staining of the mature zone in non-infected roots treated with 600nM isoxaben or 0.01% DMSO.

Supporting Information Fig. S4 DAB staining of *Heterodera schachtii* infected roots treated with 600nM isoxaben.

