## Supporting information for "WOX11-mediated cell size control in Arabidopsis attenuates fecundity of endoparasitic cyst nematodes"

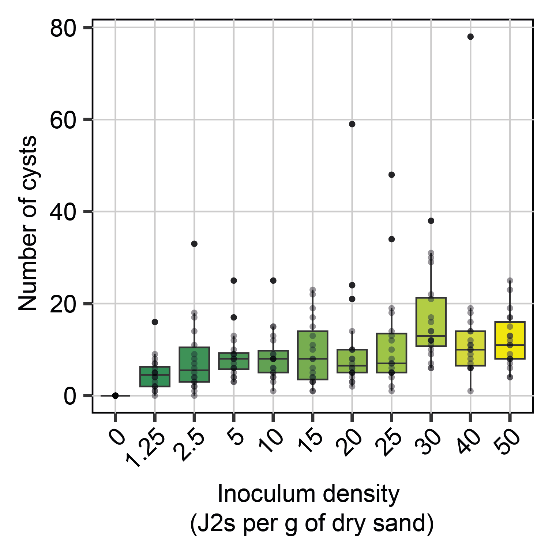

**Fig. S1** Number of *Heterodera schachtii* cysts at increasing inoculation densities in wild-type Col-0 grown in pots. 21-day-old Arabidopsis Col-0 plants were inoculated with increasing nematode densities ranging from 0 to 50 juveniles per g of dry sand. At 28 dpi, cysts were extracted and counted.

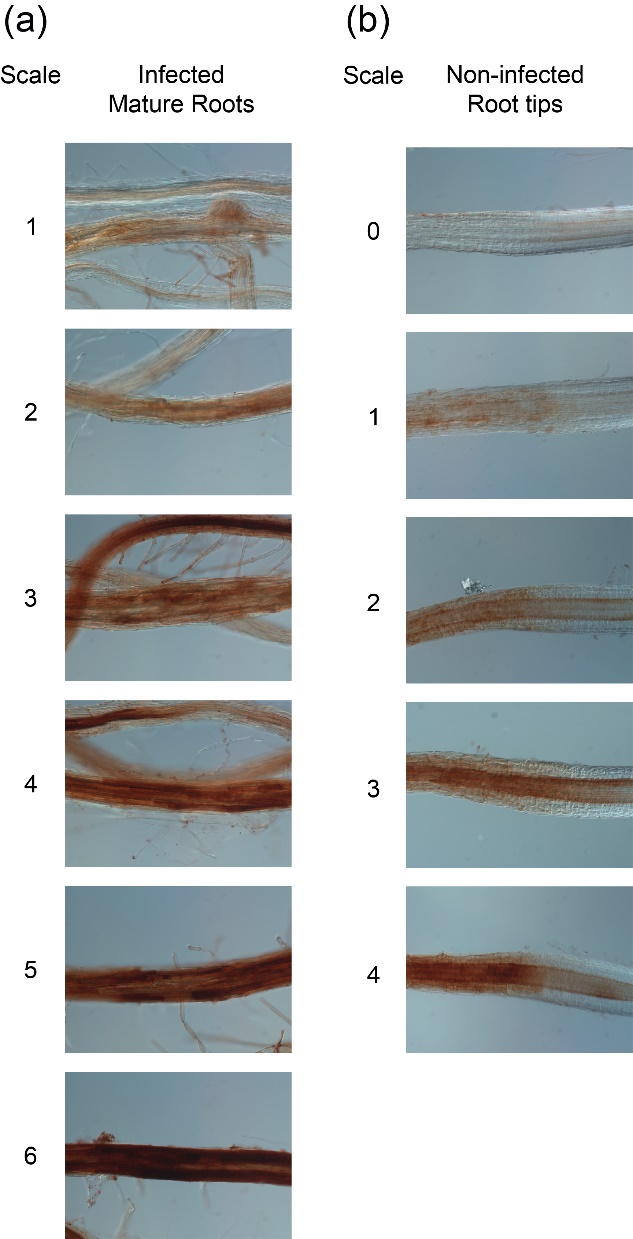

**Fig. S2** Scale used for scoring the DAB staining of Arabidopsis roots. Four-day-old Arabidopsis seedlings were inoculated with 15 *Heterodera schachtii* J2s or a mock solution. At 5 dpi, seedlings were incubated in a DAB staining solution for 2 hours in the dark. (a) DAB staining of nematode infection sites on a scale from 1 to 6. (b) DAB staining of non-infected root tips on a scale from 0 to 4.

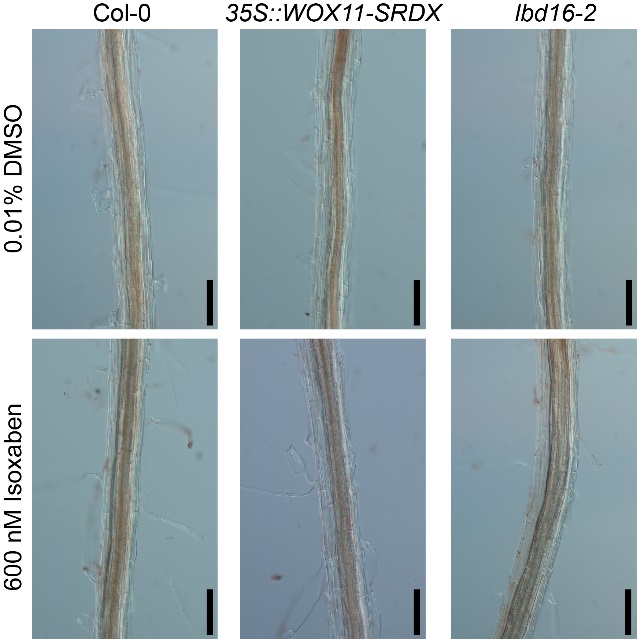

**Fig. S3** DAB staining of the mature zone in non-infected roots treated with 600nM isoxaben or 0.01% DMSO. Nine-day-old Col-0, 35S::WOX11-SRDX, and lbd16-2 Arabidopsis seedlings were transferred to liquid Knop medium containing either 600 nM isoxaben or 0.01% DMSO. After 5 hours, seedlings were incubated in a DAB staining solution for 2 hours in the dark. Scale bar is 100 μm.

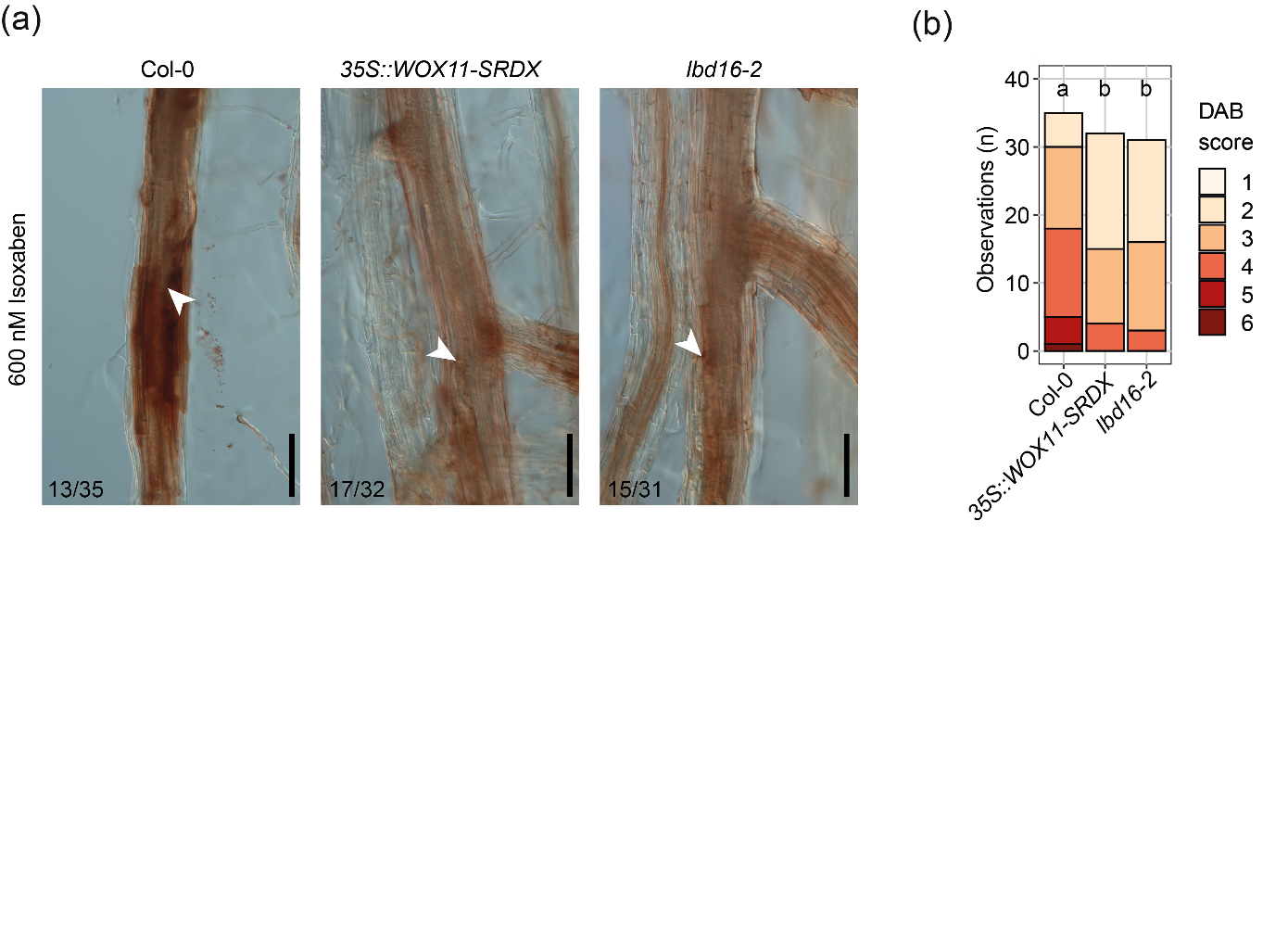

**Fig. S4** DAB staining of *Heterodera schachtii* infected roots treated with 600nM isoxaben. Four-day-old Col-0, 35S::WOX11-SRDX, and lbd16-2 Arabidopsis seedlings were inoculated with 15 *H. schachtii* J2s. At 5 dpi, seedlings were transferred to liquid Knop medium containing 600 nM isoxaben. After 5 hours, DAB staining was performed for 2 hours in the dark. (a) Representative pictures of nematode syncytia. White arrowheads indicate the nematode head. Numbers at the bottom left corner indicate how often a similar phenotype as shown in the representative pictures was observed. Scale bar is 100 μm. (b) Scoring of DAB staining intensity on a scale from 1 to 6, based on the pictures shown in Figure S2. Bar graphs indicate how often a certain DAB score is observed in each genotype. This experiment was performed three times with similar outcomes and data was pooled for statistical analysis. Significance of differences between genotypes was calculated by and Aligned Ranks Transform (ART) non-parametric ANOVA followed by an ART-contrast test for multiple comparisons (P<0.0001, n=31-35). Different letters indicate statistically different groups.
